# Keeping time and rhythm by replaying a sensory-motor engram

**DOI:** 10.1101/2022.01.03.474812

**Authors:** Victor de Lafuente, Mehrdad Jazayeri, Hugo Merchant, Otto Gracía-Garibay, Jaime Cadena-Valencia, Ana M. Malagón

## Abstract

Imagine practicing a piece of music, or a speech, solely within the mind, without any sensory input or motor output. Our ability to implement dynamic internal representations is key for successful behavior, yet how the brain achieves this is not fully understood^1–4^. Here we trained primates to perceive, and internally maintain, rhythms of different tempos and performed large-scale recordings of neuronal activity across multiple areas spanning the sensory-motor processing hierarchy. Results show that perceiving and maintaining rhythms engage multiple brain areas, including visual, parietal, premotor, prefrontal, and hippocampal regions. Each area displayed oscillatory activity that reflected the temporal and spatial characteristics of an internal metronome which flexibly encoded fast, medium, and slow tempos on a trial-by-trial basis. The presence of widespread metronome-related activity across the brain, in the absence of stimuli and overt actions, is consistent with the idea that time and rhythm are maintained by a mechanism that internally replays the stimuli and actions that define well-timed behavior.

## Introduction

Internalizing the spatiotemporal structure of external events is essential for successful interactions between organisms and their environment^5,6^. Playing music or dancing are wonderful examples of our capacity to develop internal representations of time that help to adequately synchronize our actions with events in the external world. When we track events that change within seconds, or fractions of a second, we rely on something that subjectively feels like an internal chronometer^7^. Understanding the neuronal mechanisms underpinning this internal clock is fundamental in neuroscience. We know that a wide array of brain structures are engaged when subjects perceive time and generate actions within small temporal windows^8–11^. Areas comprising the motor system consistently activate with ramping or cycling patterns of neuronal activity^12–18^, and these dynamics predict the time of past or upcoming events and actions^19–22^. Sub-cortical areas, like the hippocampus^23–28^, basal ganglia^29–33^, or the cerebellum^34–37^, have neurons that increase their activity at preferred time points and preferred interval durations, measured in either absolute (elapsed time) or relative time units (e.g. half the total elapsed time). These important previous studies show that time-perception and time-reproduction are achieved by a distributed mechanism recruiting premotor, motor, association, and sub-cortical areas to varying degrees depending on whether the behavioral task requires subjects to perceive time and/or generate timed behaviors. But what is the fundamental mechanism behind time perception and time reproduction?

To study how the brain perceives and reproduces time we used a novel behavioral task in which subjects perceived and internally maintained a visual metronome^38^. By recording from visual (V4), premotor (SMA), parietal (LIP and MIP), prefrontal (PFC), and medial temporal areas (hippocampus) while non-human primates performed the metronome task, we show that oscillatory activity encoding the metronome entrainment engages the complete hierarchy of brain areas, from sensory, to association, to motor areas. Importantly, we observed that when the metronome was no longer available externally, but subjects were required to keep its tempo, each of the recorded areas maintained patterns of oscillatory activity that encoded the tempo and spatial location of the metronome. These results support the view that time perception and time reproduction are achieved by internally replaying the entire repertoire of sensory, motor, and cognitive representations that are engaged for precise spatiotemporal coordination of actions with external events.

## Results

### Subjects perceive and maintain rhythms of varying tempos

We trained two rhesus monkeys to perform a metronome task in which they had to perceive, and internally maintain, a temporal rhythm defined by a visual stimulus that periodically alternated left and right across a central eye fixation point on a screen (Fig. 1a, Supplementary Methods; all methods for this paper are provided in the Supplementary Information). The task initiated by visually presenting the alternating metronome (3 intervals, *entrainment* epoch), then the metronome disappeared, and the animals had to internally estimate its position as a function of time. At the middle of a randomly selected interval within the *maintenance* epoch (1 to 6 intervals; uniform distribution), a *go-cue* instructed the subjects to reveal their estimated location of the metronome by touching the left or right side of a touchscreen (*left* or *right* behavioral choices). It is important to note that there are no stimuli, and no motor actions are required during the *maintenance* epoch before the *go-cue*. This allowed us to study how time is dynamically encoded and how tempos of different rhythms are internally maintained in the absence of sensory or motor cues.

**Fig 1.**
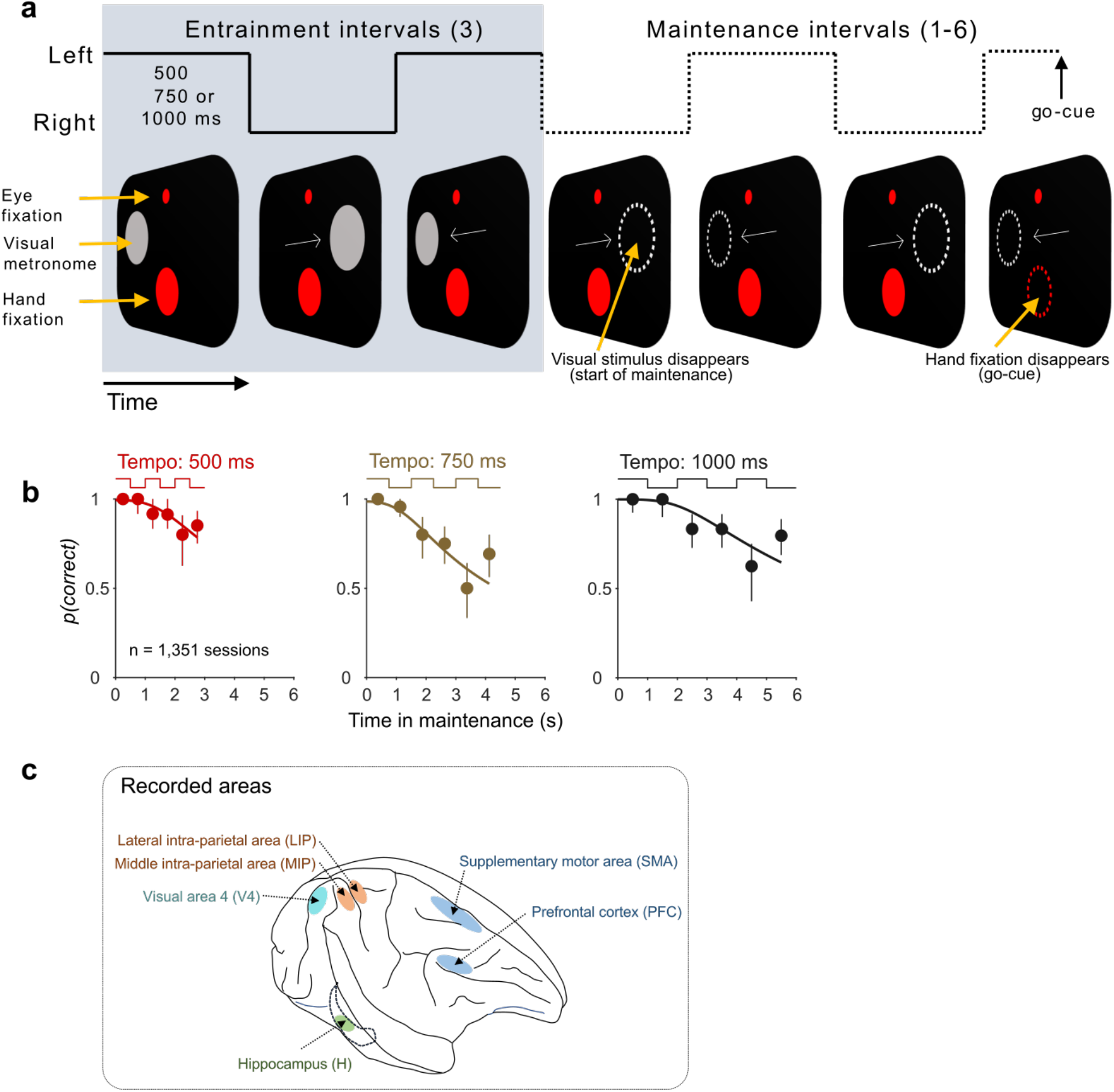
Monkeys performed the metronome task while recordings were made from six brain areas. **(a)** In each trial, the animal was presented with a visual metronome (gray circle) alternating between left and right with a fixed tempo (entrainment intervals). The tempo for each trial was chosen randomly to be 500, 750, or 1000 ms. After the entrainment intervals, the animals had to maintain the tempo for up to six additional intervals (maintenance intervals; four are illustrated for a left-starting metronome trial). Eye and hand fixation were required throughout the trial, up to the go-cue that instructed the animals to move their hand from the hand fixation point to the estimated location of the metronome at the time of the go cue. **(b)** The probability of a correct response p(correct) is plotted as a function of elapsed time in the maintenance epoch (mean ± IQR). As expected, p(correct) decreases with time indicating that the internal metronome increasingly deviates from the correct tempo during maintenance. Solid lines are fits from a model making use of the scalar property of timing in the context of the metronome task^38^ (1,351 sessions were pooled to generate the behavioral curves; 955 monkey I; 396 monkey M; Extended Data Fig. 1). **(c)** We recorded from six brain areas (independently) including the frontal lobe (SMA, PFC; blue), parietal lobe (LIP, MIP; orange), occipital lobe (V4; cyan), and temporal lobe (Hippocampus; green; Extended Data Fig. 1).

Subjects adequately performed the metronome task, estimating and maintaining metronomes with fast, medium, and slow tempos that varied randomly across trials^38^. Subjects made few mistakes when they had to indicate the metronome’s position early in the *maintenance* epoch (96.5 ± 0.2% correct responses for the first *maintenance* interval; mean ± s.e.m. across the 3 tempos; Fig. 1b, Extended Data Fig. 1), and the proportion of correct responses decreased as a function of elapsed time. This behavior is consistent with the scalar property of time perception^39,40^ predicting that variability in the estimation and maintenance of the metronome will result in increasingly larger deviations from the correct tempo as time elapses.

### Single neurons and network activity oscillate at the tempo of the internal metronome

While subjects performed the metronome task, we recorded neurons in visual area 4 (V4), in the lateral and medial cortices of the intra-parietal sulcus (LIP and MIP, respectively), in the supplementary motor area (SMA), in the prefrontal cortex (PFC), and in the Hippocampus (Fig. 1c; separate recording sessions; Supplementary Methods). Example neurons from each area illustrate two main findings: (1) Neurons are responsive to the visual presentation of the metronome, and they also show oscillations during *maintenance*, an epoch where the metronome is no longer visible. (2) This oscillatory activity stretches to match the tempo of the metronome, which varied randomly on a trial-by-trial basis (Fig. 2).

**Fig 2.**
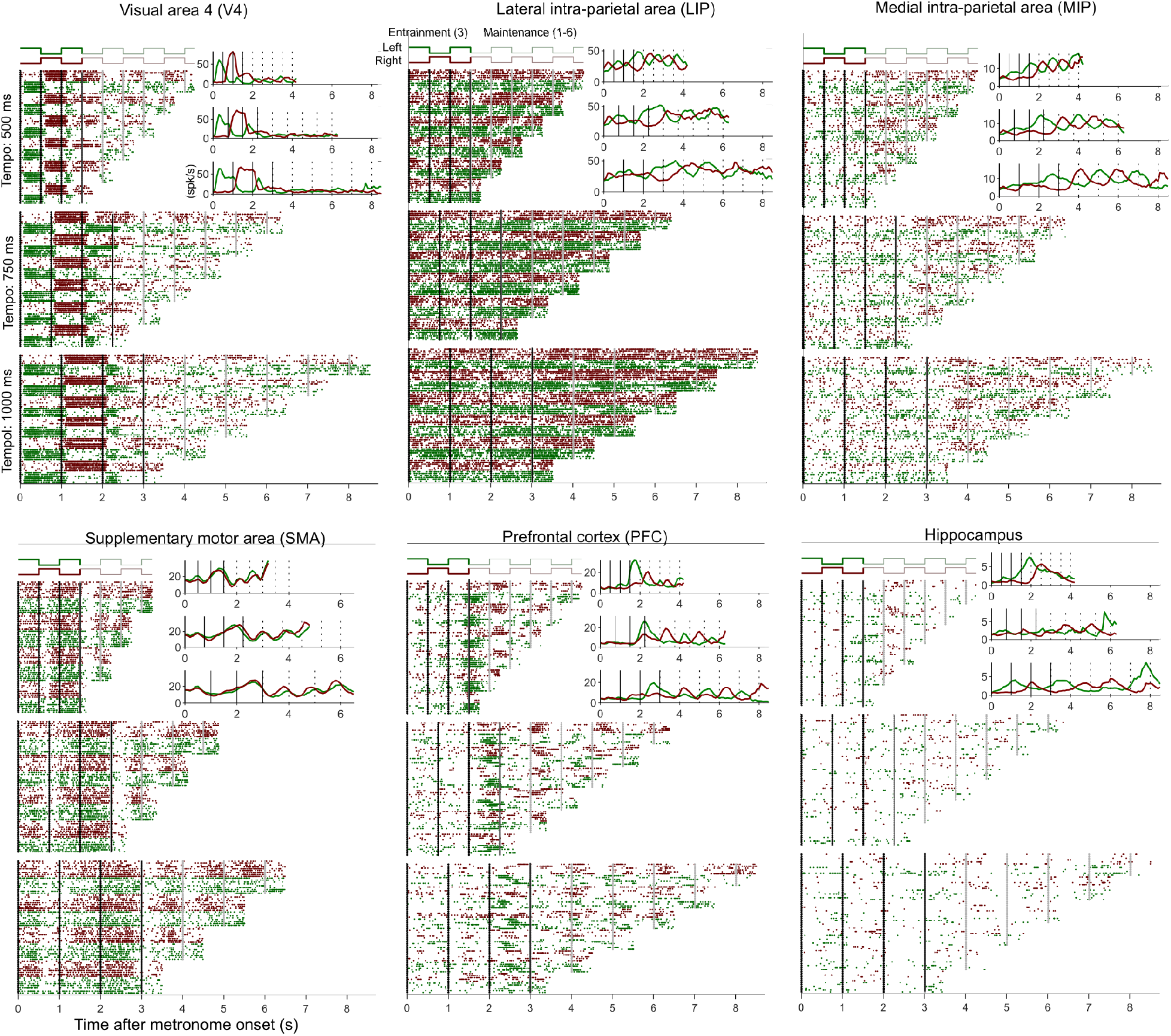
Spiking activity of example neurons from each recorded area. Trials are aligned to the start of the visual stimulus initiating the metronome. Each dot marks the time of a spike; green and red dots represent left- and right-staring metronomes, respectively. The metronome’s position is illustrated by the traces at the top. Trials are sorted by tempo (interval durations of 500, 750, 1000 ms), location (left-right), and number of maintenance intervals presented (1-6; 1-4 for the SMA). Insets in each rasterplot depict mean firing rates (spk/s) for left and right-starting metronomes (green and red lines, respectively), for the three tempos. Prominent oscillatory activity that scales with the metronome’s tempo is observed in each example neuron.

The mean firing rate across recorded neurons shows that visual area V4 responded with a short latency to the onset of the visual stimuli within the neuron’s preferred location (Fig. 3a, V4; Supplementary Methods). Importantly, V4 neurons continued to oscillate when the stimulus was no longer visible during the *maintenance* epoch. The rhythmic activation of V4 in the absence of a visual stimulus suggests that spatial attention might be alternating between the left and right visual fields as subjects internally track the position of the stimulus. We speculate that rhythmic activation of this visual sensory area might be an indication that the monkeys generate an internal dynamic visual representation of stimuli that is no longer visible but whose position is still relevant for a correct behavioral outcome^41,42^. Parietal area LIP also displayed strong sensory responses and, like parietal area MIP, maintained these oscillatory responses when the stimulus was no longer visible (*maintenance* epoch; Fig. 3a; LIP). The know role of the supplementary motor area (SMA) in the preparation of future movements is well illustrated in our data as patterns of oscillatory activity mounted over increasing firing rates anticipating the end of the trial (Fig. 3a; SMA). We found that PFC and the hippocampus also contained neurons that displayed oscillatory patterns during the *entrainment* and *maintenance* epochs of the metronome task (Fig. 3a, PFC, Hippocampus). Note that all oscillatory signals during the *maintenance* epoch play out in the absence of any external stimulus or overt movements. Thus, our results strongly suggest that the fast (500 ms), medium (750 ms), and slow (1000 ms) oscillatory patterns reflect the encoding of the tempos of internally maintained metronomes.

**Fig 3.**
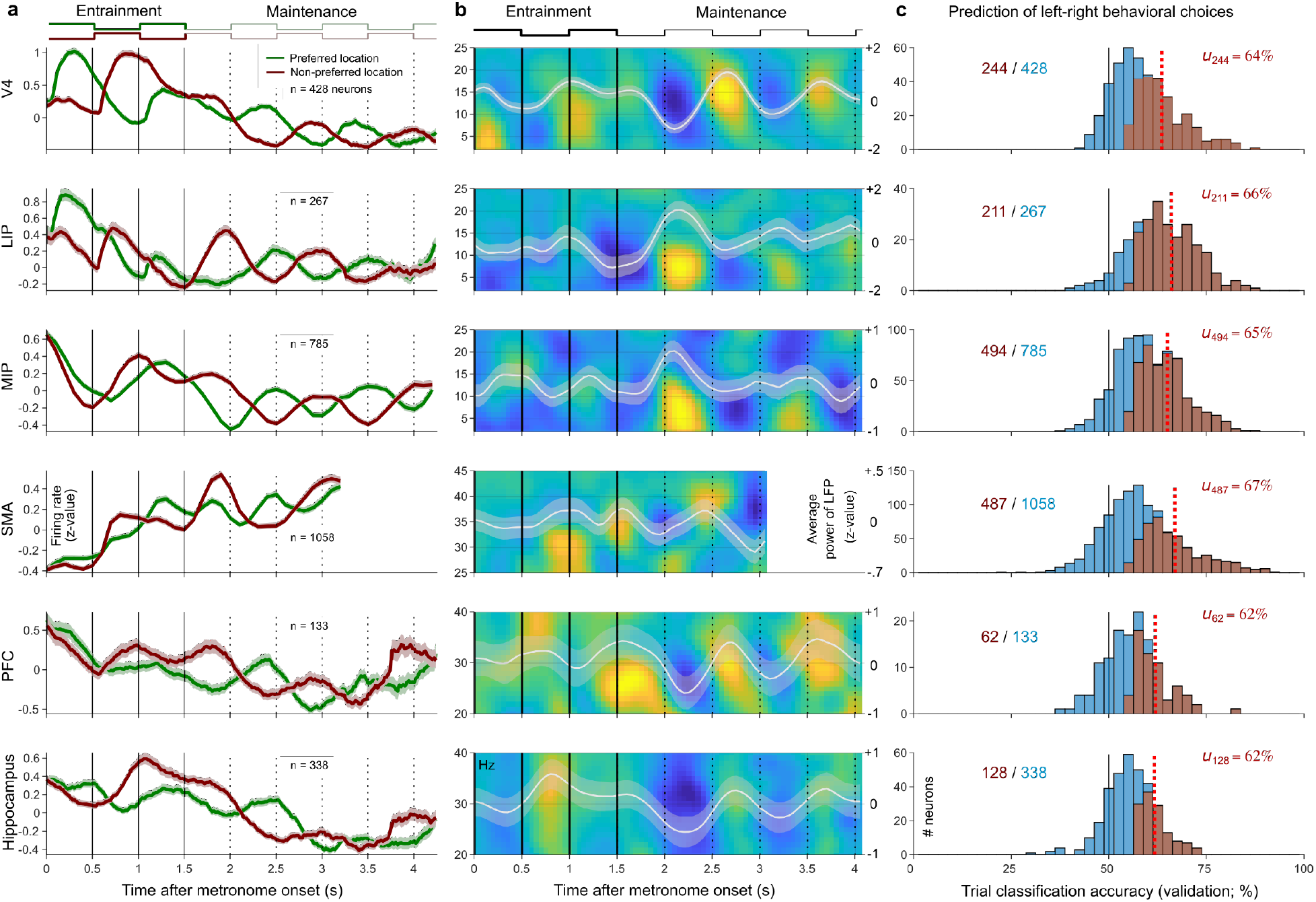
Activity of neurons and field potentials show oscillations that are predictive of behavioral choices. **(a)** Mean firing rates across recorded areas (z-values; mean ± s.e.m). Note that each area displayed prominent oscillations in the entrainment and maintenance epochs of the metronome (500 ms tempo shown). **(b)** Spectral power of the local field potentials (LFPs) during the metronome task. Color shows the power difference between right- and left-starting metronome trials. White lines indicate this difference across frequencies (z-values; mean ± s.e.m). Note that alternating yellow and blue colors correlate with changes in the metronome’s position, even in the maintenance epoch when the stimulus is no longer visible. **(c)** Classification accuracy of behavioral choices using the spiking of single neurons (SVM classifier). The numbers of significant neurons are in red (p<0.05), and red dotted lines indicate their mean accuracy. Black line at 50% indicates chance accuracy.

In addition to spiking activity from single neurons, we also analyzed power changes in the local field potential (LFP), a signal that is known to reflect large-scale network activity. Results indicate that population of neurons (likely including circuits of principal neurons and interneurons), were engaged in the *entrainment* and *maintenance* epochs of the metronome (Fig. 3b). Remarkably, LFP power within a wide-band set of frequencies oscillated in synchrony with the tempo of the metronome, both in the *entrainment* and *maintenance* epochs (Fig. 3c. see Extended Data Fig. 2a for a larger set of frequencies). Notably, for each recorded area the oscillatory power of LFP was stronger when the stimulus was no longer visible (p<0.01 for all areas; Extended Data Fig. 2b; Supplementary Methods). This suggests an increase in sub-threshold and local processing demands in the *maintenance* epoch.

To test whether the oscillatory activity of neurons was indeed related to tracking the metronome’s position, we used a support-vector machine (SVM) classifier. The classifier estimated the accuracy with which *left-right* behavioral choices could be predicted from the activity of single trials (single neurons) during the *maintenance* epoch. The cross-validated results show that mean prediction accuracy in each recorded area was significantly beyond 50%, with a large proportion of neurons that individually predicted behavioral choices (Fig. 3c; Supplementary Information). Thus, even when the metronome is no longer visible, and no movements are being made, each area contains significant information to decode the internal metronome guiding behavior (Fig. 3c; both correct and incorrect choices could be decoded; Extended Data Fig. 2c).

### Encoding the metronome’s tempo and location

To gain insight into the role each area plays in the encoding of the metronome we analyzed population activity by means of principal component analysis (PCA). This allowed us to visualize the effect that spatial location (*left*-*right*) and tempo (500, 750, 1000 ms) have on the overall dynamics of the population activity (Fig. 4; Extended Data Fig. 3). The population activity spanned by the first three PCs shows that V4 neurons do not encode the speed of the metronome. This is demonstrated by the proximity of the trajectories for the different tempos (Fig 4a, left panel). That the activity elicited by different tempos follows similar trajectories indicates that, although V4 strongly responds to the visual stimulus, those responses do not carry information about the speed of the metronome. In contrast, PCs of V4 activity clearly distinguish the spatial location of the metronome. This is illustrated by the distinct trajectories that the population follows in response to *left-starting* and the *right-starting* metronomes (Fig. 4a, right panel; red vs green lines).

**Fig 4.**
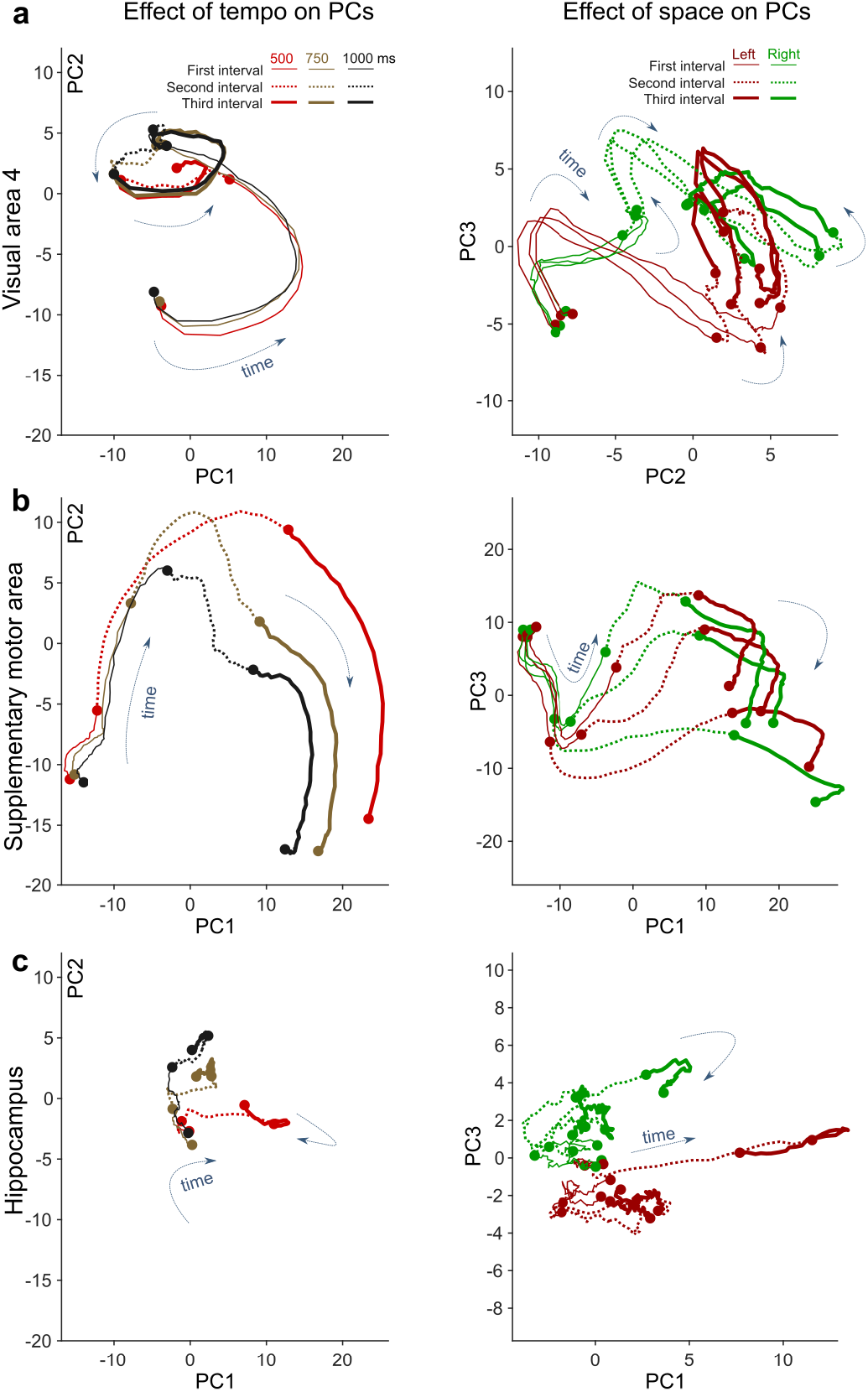
Dynamics of neuronal populations in V4, SMA, and Hippocampus in the entrainment of the metronome. **(a)** The first two principal components (PC1, PC2) in area V4 for different tempos are plotted in the left panel. Population activity for different tempos (500, 750, 1000 ms) evolve close to each other, illustrating that V4 activity does not distinguish fast, medium, or slow tempos. The right panel shows that left- and right-starting metronomes follow different trajectories, thus demonstrating spatial selectivity. **(b)** Population activity in SMA follow separate paths for the metronomes of different tempos thus illustrating the capacity of SMA to encode the metronome’s tempo (left panel). Conversely, the right panel shows that trajectories for left- and right-starting have similar dynamics, demonstrating that population activity in SMA has weak spatial selectivity. **(c)** The metronome’s tempo is weakly encoded in the hippocampus (left panel). However, trajectories for left- and right-starting metronomes do not cross or overlap, indicating that this area is able to encode the spatial location where the metronome started (right panel).

Contrasting with V4 neurons, SMA population activity follows distinct trajectories for the different tempos of the metronome, particularly during the last two entrainment intervals (Fig. 4b; left panel). This result indicates that SMA encodes the speed at which the visual stimulus alternates on the screen. Also in contrast to V4 neurons, the activity of the SMA follows similar trajectories for *left-starting* and *right-starting* metronomes, illustrating that this area does not strongly encode the spatial location of the metronome (Fig. 4b left panel; red vs green lines).

The hippocampus is an interesting structure to contrast with the SMA and V4 because it is not motor, nor sensory, nor related to sensory-motor transformations as the parietal and prefrontal cortices are known to be^43^. Although different trajectories for different tempos are observed in the hippocampus, the change in activity is much less pronounced than that observed in V4 and SMA (Fig. 4c, left panel; firing rates were normalized across neurons and areas to avoid this potential confound). Interestingly, the hippocampus shows a unique pattern of activity that was not observed elsewhere in our recordings: *left-starting* and *right-starting* metronomes give rise to separate trajectories that remain separate for the duration of the *entrainment* epoch (Fig. 4c, right panel). This is a remarkable observation because keeping track of where the metronome started is not useful to solve the task (where the metronome starts is uncorrelated to where it will be at *go-cue* time). This pattern of activity is consistent with the hippocampus’ proposed role as a generator of sequential state spaces that gives context and temporal order to externally and internally generated brain states^27,44^.

To quantify the observations made with PCA we performed a demixed-PCA analysis (dPCA) that allowed us to measure the amount of information each area encodes, explicitly in relation to (1) the spatial location (*left*-*right*), (2) the tempo (500, 750, 1000 ms), and (3) the total elapsed time within the trial. The results corroborate the PCA observations by demonstrating that V4 neurons strongly encode space, while the SMA shows larger encoding weights for tempo (Fig. 5a). By performing this analysis for each area, and plotting the spatial vs temporal information, we were able to identify a distributed hierarchy of neural representations for spatial and temporal properties of the underlying metronome (Fig. 5b, Extended Data Fig. 5). Results show that V4 and LIP have the largest encoding capacity for the spatial location of the metronome, while SMA displays the largest capacity for tempo encoding, followed by the parietal areas LIP and MIP (Fig. 5b). In addition to the temporal rhythm of the metronome, dPCA estimated how much the activity of each area varies in relation to the passage of total elapsed time. The results show that the ability of an area to encode the metronome’s tempo is highly correlated with its capacity to encode total elapsed time (Fig. 5c). This finding remarkably illustrates that time and rhythm perception share the same fundamental neuronal mechanisms. We note that the capacity of areas to encode space and time is inversely correlated with their *dimensionality* (measured as the number of PCs needed to explain most of the activity variance) (Fig. 5d). Areas V4 and SMA need only a third of the PCs needed to explain 80% of population activity variance as compared to the hippocampus. We speculate that the low dimensionality of areas encoding space and time would make it straightforward for the rest of the brain to perform a readout of those parameters, conveying flexibility and capacity for generalization of the metronome to new behavioral requirements.

**Fig 5.**
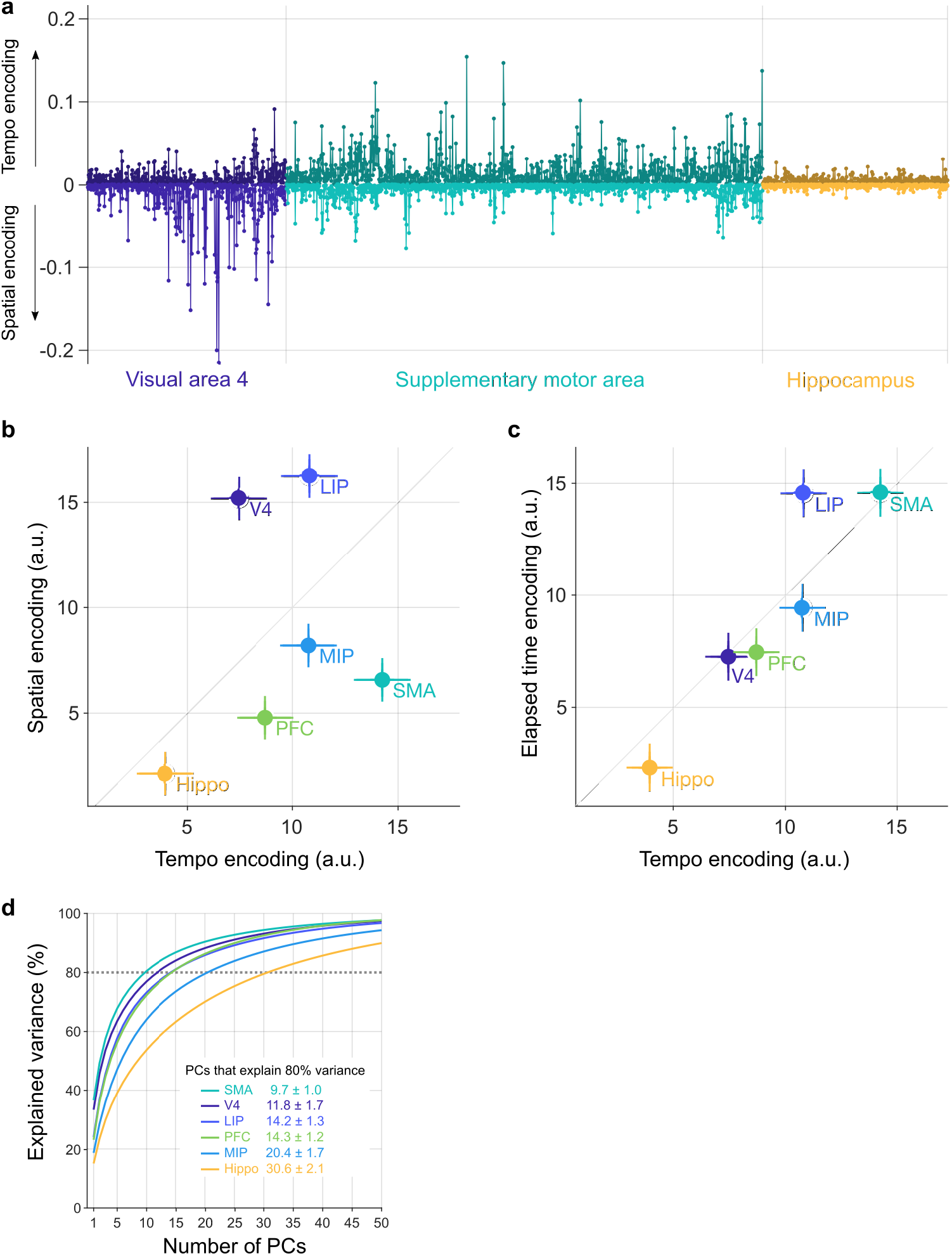
Encoding the tempo and spatial location of the metronome. **(a)** The capacity of each neuron (x-axis) to encode tempo and space (y-axis; encoding weights) was calculated using dPCA and is shown for areas V4, SMA, and Hippocampus (Extended Data Fig. 4 plots all neurons for each area). Note the strong spatial selectivity of V4 neurons and the strong tempo encoding of SMA neurons. **(b)** Plotting dPCA weights (mean ± s.e.m) for spatial and tempo information reveals a hierarchy of brain areas in relation to their capacity to encode these metronome characteristics. Note that LIP has the largest weights for spatial encoding, and that the SMA has the largest weights for tempo encoding. **(c)** The capacity to encode the metronome’s tempo (x-axis) is highly correlated with the capacity to encode total elapsed time (y-axis). Note that the fronto-parietal circuit comprising LIP and SMA excels at encoding tempo and elapsed time. **(d)** The cumulative percent of explained variance is plotted as a function of the number of PCs. The inset table shows the number of PCs required to explain 80% of activity variance across areas (mean ± s.d.; bootstrap). Note that areas SMA and hippocampus are at the extremes of a hierarchy of activity dimensionality.

## Discussion

We found that frontal, parietal, occipital, and hippocampal regions of the brain display oscillatory activation patterns reflecting the perception of temporal rhythms that are internally maintained to solve a metronome task. We show that the brain flexibly adapts the tempo of its internal metronome, on a trial-by-trial basis, with the underlying neuronal oscillations scaling their pace accordingly. Importantly, we demonstrate that the spatial and temporal characteristics of the metronome are maintained in the absence of sensory stimuli and movements. During this internal representation of the metronome, all recorded areas contained information that significantly predicted the subjects’ behavioral choices.

While it would be possible to provide an *ad hoc* explanation for the involvement of each recorded area in the metronome task (e.g., a sensory role for V4, motor preparation for the SMA, spatial attention for the parietal cortices, and cognitive control for the PFC), a more parsimonious account of the data might be that the brain solves the metronome task by performing an internal replay of the entire repertoire of sensory and motor events associated with the task. Accordingly, we hypothesize that the internal representation of structured spatiotemporal events (engrams, such as the visual metronome) can be explained by the internal replay of the sensory, motor, and cognitive activity patterns generated in the brain while learning the task. Three main observations support the engram replay hypothesis: (1) A sensory visual area (V4), and its related parietal association cortices (LIP, MIP), dynamically keep track of the metronome position across time in the absence of a visual stimulus. (2) A premotor area (SMA) shows oscillatory activity during the *entrainment* epoch, where no movement is ever required. This observation suggests that subjects might be *internally reaching* for the stimulus as a proxy to estimate the metronome’s tempo. (3) The hippocampal activity keeps track of the metronome’s starting position even when this information does not contribute to solving the task. This observation is consistent with the idea that the hippocampus might provide a general context for other areas to correctly replay the metronome’s engram. We note that our findings, far from disagreeing with previous results, encompass them into a unifying framework in which the cognitive process that we call *timing* is achieved by making use of the brain’s known machinery for constructing internal models of the past, present, and future states of the word^45–48^. We showed that estimating and maintaining a metronome engages a large distributed sensory-motor engram. These areas were found to be organized in a hierarchy with respect to temporal structure, with sensory areas representing the current state of the world, the temporal lobes maintaining past information, and the frontal and parietal lobes anticipating upcoming changes.

## Data availability

Data is available from the corresponding author upon reasonable request.

## Code availability

The analyses were performed in MATLAB (Mathworks) and code is available from the corresponding author upon reasonable request.

## Acknowledgements

We thank Edgar Bolaños for technical assistance, Juan Ortiz and Pamela García for obtaining the MRI images, and members of the Lafuente lab for fruitful comments. We thank Jessica Norris for proofreading the manuscript. This work was supported by grants from CONACYT, PAPIIT (IG200521), and the Massachusetts Institute of Technology (MIT) International Science and Technology initiative.

## Author contributions

V.d.L and M.J. designed the experiments. V.d.L., O.G.G., J.C.V., and A.M.M trained the monkeys and performed the recordings. V.d.L. wrote the initial draft of the manuscript. V.d.L. analyzed the data. All authors contributed to the scientific content of the work.

**Correspondence and requests for materials** should be addressed to V.d.L.

## Extended Data

**Extended Data Fig. 1.**
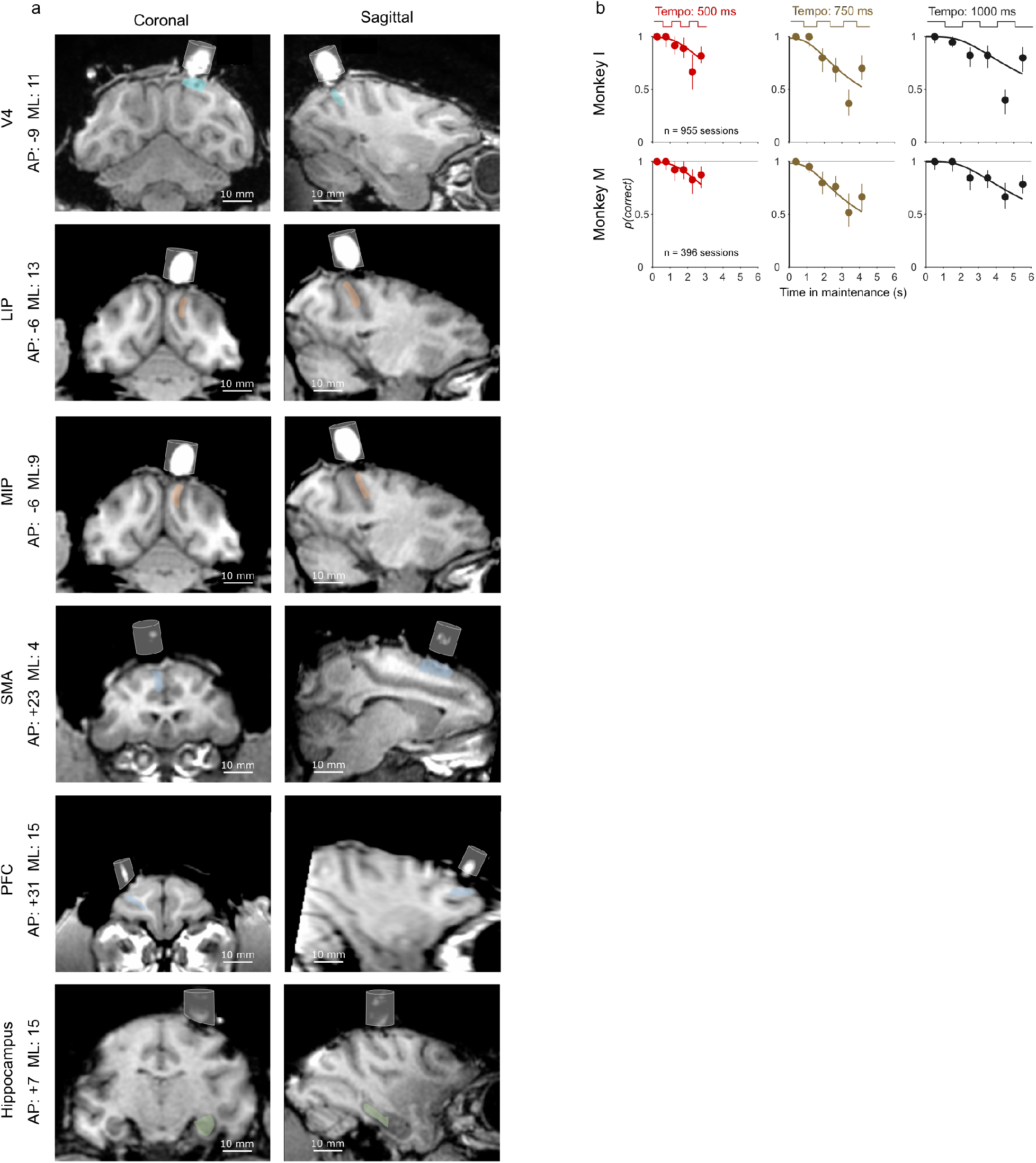
Recording locations and behavioral performance. **(a)** Magnetic resonance images of the recording locations. Antero-posterior (AP) and medio-lateral (ML) stereotaxic coordinates (ear-bar zero) are given for the center of the recording chambers (white cylindrical area) for each area. Colored areas indicate the estimated recording locations within the brain. **(b)** Behavioral performance in the metronome task for monkey I (upper row) and monkey M (lower row). The proportion of correct responses *p(correct)* is plotted as a function of elapsed time in the *maintenance* epoch. Conventions are the same as in main Fig 1b. Note how both monkeys make more mistakes when locating the metronome after 5 maintenance intervals. This bias might be explained by a tendency to choose the location of the last visible stimulus after a long maintenance period, when uncertainty about the metronome’s position is greater.

**Extended Data Fig 2.**
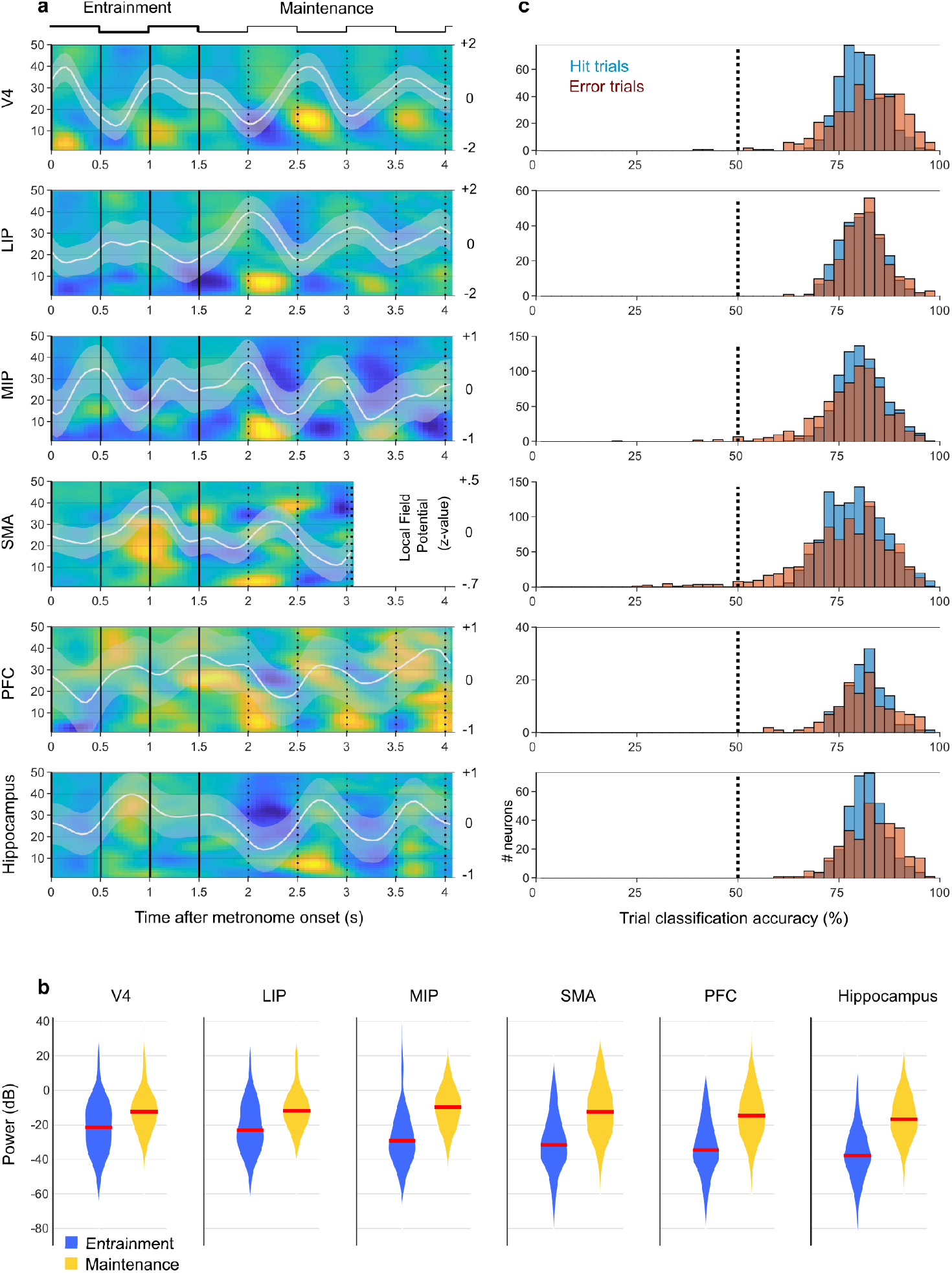
Spectral power of LFPs and accuracy of classification of hit and error trials. **(a)** Spectral power of the local field potentials (LFPs) during the metronome task across the set of 1-50 Hz frequencies. Color indicates the power difference between *right-* and *left-starting* metronome trials. The white line indicates the spectral power across the displayed frequencies (z-values; mean ± s.e.m). Conventions are the same as in main Fig 3c. Note the alternating yellow and blue colors that correlate with the changes in the metronome position, even in the maintenance epoch when the metronome is no longer visible. **(b)** Violin plots comparing the power of oscillations in the LFPs during the *entrainment* and *maintenance* epochs. In all areas oscillations of the LFP power were higher during *maintenance*, an epoch in which the stimulus is no longer visible (p<0.01 for all areas; paired t-tests; red line indicates the median oscillatory power across recording sessions). **(c)** Single trial classification accuracy of behavioral choices. Black dotted line at 50% indicates chance accuracy. Blue bars indicate the accuracy with which correct (hit) trials could be classified. Red bars indicate accuracy for the incorrect behavioral choices (error trials). Note that neurons in each recorded area contained enough information to classify a large proportion (~80) of hit and error trials (the full SVM model was used, *i.e.* all trials contributed to training the model, hence the high classification accuracy).

**Extended Data Fig 3.**
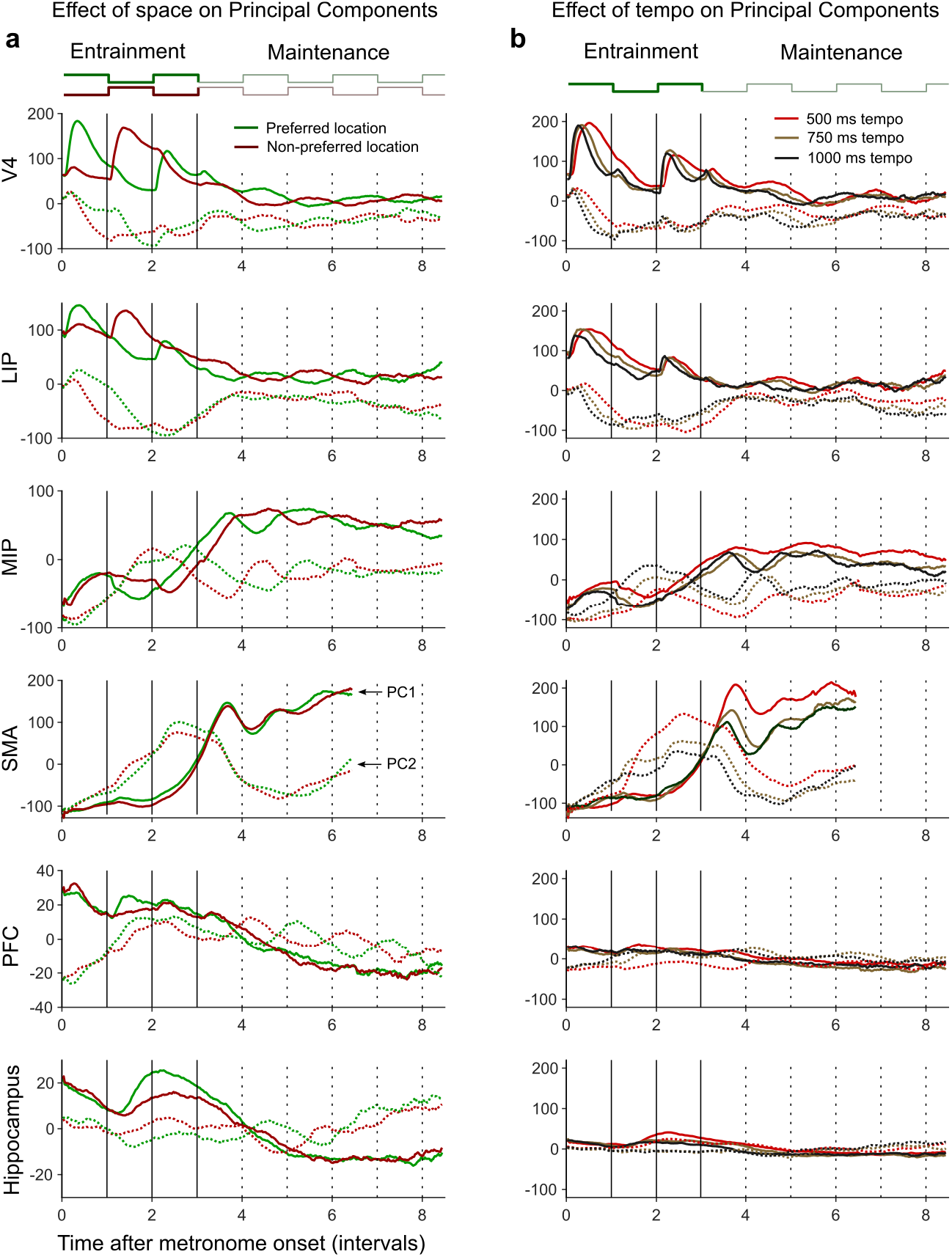
Effect of space and tempo on the dynamics of the first two PCs. **(a)** The panels plot the first two PCs of population activity for each area (PC1, continuous lines; PC2, dotted lines). Green color illustrates PC activity when the metronome started at the neurons’ preferred location. Red color shows the activity elicited by the metronome starting in the non-preferred location. Note that V4, and parietal areas MIP and LIP, have activity with strong spatial preference that extends into the *maintenance* epoch. Area SMA does not show strong spatial preference. The PFC and hippocampus both show oscillatory activity, although with smaller intensity (note the differences in the y-axis scales). **(b)** The panels show the first two PCs of population activity in response to metronomes of different tempos (500, 750, 1000 ms; plotted on a common time axis in interval units). Note how areas V4 and LIP have largely overlapping dynamics for the different tempos, suggesting that other areas might be responsible setting and maintaining the metronome’s tempo. Areas MIP and SMA display population activity that follow separate trajectories for different tempos. Note that in these panels *y*-axis scale is the same for all areas.

**Extended Data Fig 4.**
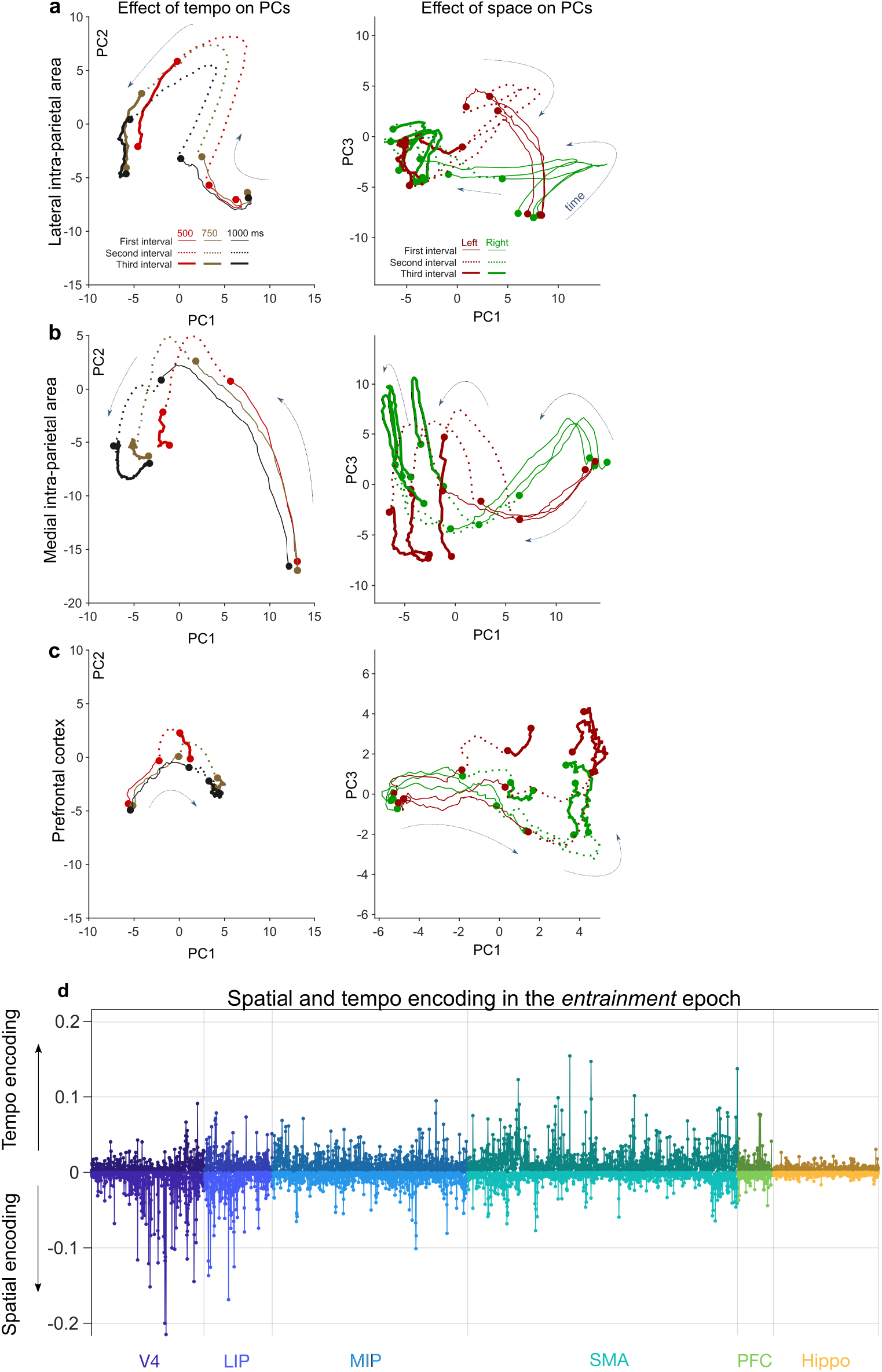
Dynamics of neuronal populations in LIP, MIP, and PFC in the *entrainment* phase of the metronome. **(a)** The first two principal components (PC) in area LIP for different tempos (500, 750, 1000 ms) are plotted in the upper panel. Note how trajectories in the second interval are separated for different tempos (dotted lines) demonstrating that population activity in LIP, to some extent, differentiates slow, medium, and fast metronomes. The lower panel shows LIP’s spatial selectivity, illustrated by the different trajectories followed by *left-* and *right-staring* metronomes. **(b)** MIP is similar to LIP in the sense that it is sensitive to different metronome tempos (different trajectories for different tempos; upper panel) and also sensitive for spatial location (different trajectories for *left-* and *right-starting* metronomes; lower panel). **(c)** Metronome tempos in the PFC are weakly encoded as illustrated by the proximity of the trajectories (500, 750, 1000 ms; upper panel). Some space selectivity is present in PFC as illustrated by the different trajectories followed by left-starting and right-starting metronomes (lower panel). **(d)** Space and tempo encoding values for each recorded neuron (dPCA weights), grouped by recording area (colors and labels on the *x*-axis). V4 and LIP show large spatial encoding, MIP has both medium spatial and temporal encoding. SMA shows large capacity for tempo encoding. PFC and Hippocampus have modest temporal and spatial encoding capacity.

**Extended Data Fig 5.**
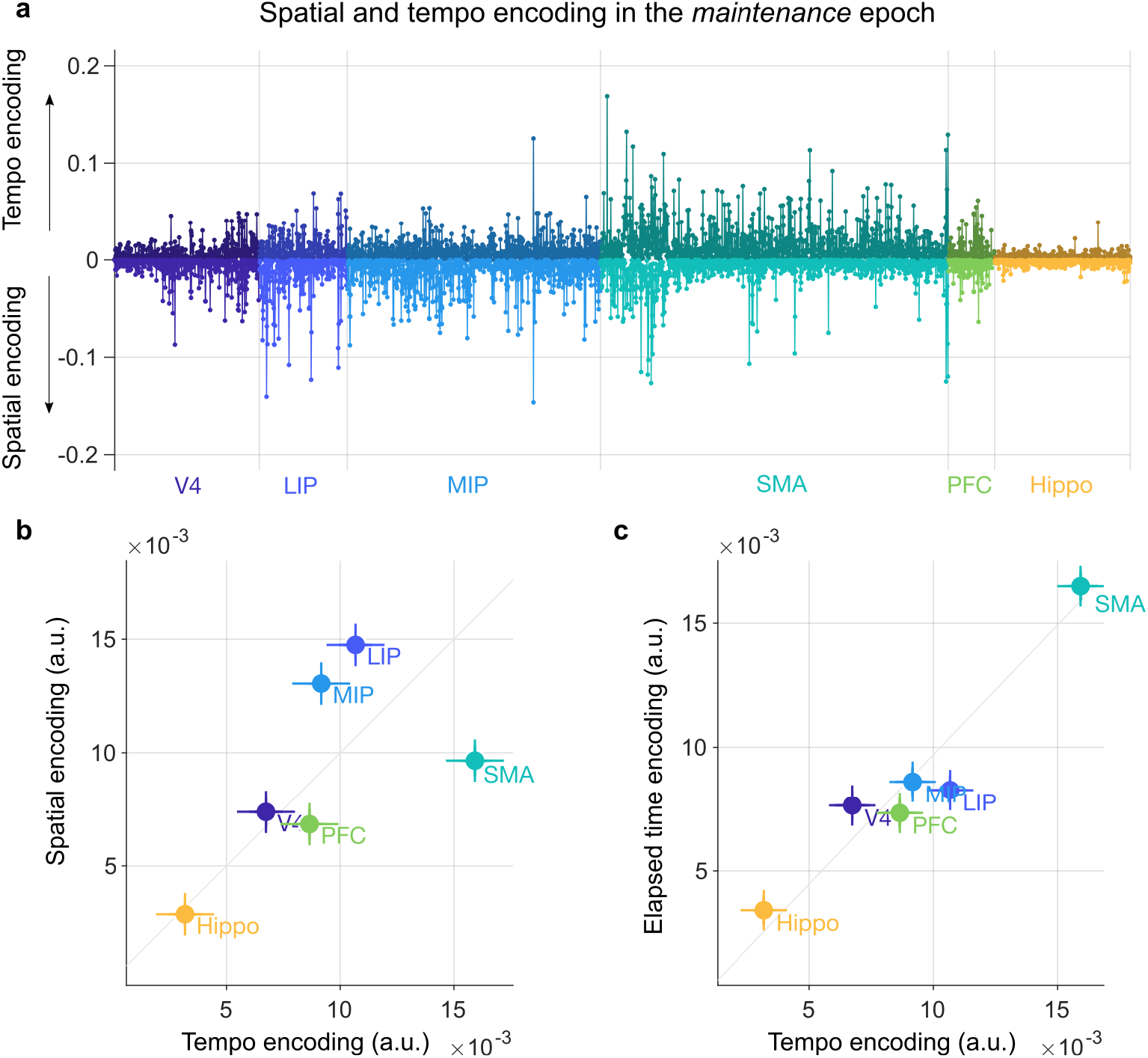
Neural encoding of space and tempo in *maintenance* epoch of the metronome task. **(a)** Space and tempo encoding values for each recorded neuron in the *maintenance* epoch of the task (grouped by recoding area). The results are similar to those obtained for the *entrainment* phase (Fig. 5a). **(b)** As compared to entrainment (Fig. 5b), in the *maintenance* epoch, V4 reduces, and MIP increases its capacity to encode the spatial information of the metronome. **(c)** The SMA retains a large capacity for tempo and elapsed time encoding, as compared with the rest of the areas that do not strongly encode total elapsed time.

## Supplementary information

### Methods

#### 1. Subjects

Two rhesus monkeys (*Macaca mulatta*) participated in this study, aged 9 (monkey I, weight: 8-9 kg, age: 9 years; monkey M, weight: 10-12 kg, age: 10 years). Experimental procedures were approved by the Ethics in Research Committee of the Institute of Neurobiology of the National Autonomous University of Mexico and agree with the principles outlined in the Guide for Care and Use of Laboratory Animals (National Institutes of Health). Monkeys were implanted with titanium head bolts and recording chambers located over the areas of interest. Placement of the chambers was guided by stereotactic coordinates and structural magnetic resonance imagining (see Extended Data Fig. 1b for the locations of the recording chambers).

#### 2. The metronome task

We have described the metronome task in detail elsewhere^1,2^. Briefly, the monkeys observed a visual stimulus (gray circle, 10° diameter, 25° eccentricity) that periodically alternated left and right from a central eye fixation point. The duration of the stimulus on each location (interval duration) was chosen pseudo-randomly from 500, 750, and 1000 ms. Subjects observed 3 *entrainment* intervals (Figure 1a), then, the visual stimulus disappeared, and the monkeys’ task was to covertly keep track of its position (left or right) as a function of elapsed time (*maintenance* epoch). After 1-6 *maintenance* intervals (pseudo-randomly chosen), a go-cue (disappearance of the hand fixation area; Figure 1a) instructed the monkeys to touch the metronome’s estimated location. Importantly, this was not an interception task since stimulus’ position did not further change after the go-cue. The go-cue was always given at the middle of one of the 1-6 *maintenance* intervals.

Subjects had to maintain eye fixation over the complete trial and hand fixation until the go-cue signal (eye tracker: ASL Eye-Track 6; touchscreen: Elo Touch 1939L). Experimental control and stimulus presentation were achieved with EXPO (developed by Peter Lennie and Rod Dotson, Center for Neural Science at New York University, New York, NY; https://sites.google.com/a/nyu.edu/expo/).

#### 3. Behavioral analysis

Performance on the metronome task was measured by psychometric curves that plotted the proportion of correct responses *p(correct)* as a function of elapsed time in the *maintenance* epoch. Fig. 1b shows one psychometric curve for each metronome’s tempo (mean ± IQR). The continuous lines are fits of a timing model which implements the scalar property of timing (variability of time estimates increases in proportion to total elapsed time^2^) in the context of the metronome task. The psychometric curves display the expected decline in correct responses as a function of elapsed time, and this due to the internal metronome increasingly getting out of synch from the correct tempo (see Extended Data Fig.1a for the psychometric curves of each monkey).

#### 4. Neuronal recordings

Recordings of extracellular spikes and local field potentials (LFPs) were performed with an array of 7 independently movable platinum-tungsten electrodes with quartz insulation (2-3 MΩ; Thomas Recording; Giessen, Germany). For the hippocampus we used an acute guide tube that vertically guided a single recording electrode down to 1 cm dorsal to the hippocampus. The remaining 1 cm was traveled by the electrode alone. Spike activity was sampled at 30 kHz and single units were isolated online (Cerebus acquisition system, Blackrock Microsystems; Salt Lake City, UT, USA). LFPs were obtained by band-pass filtering the electrode signal at 0.5-500 Hz, at a 1Khz sampling rate. Offline, the LFP signal was band-pass filtered to the 1-50 Hz band. Neuronal recordings were obtained from the two monkeys in all areas, except for the prefrontal cortex and the hippocampus, that were recorded only in monkey I. All neurons that remained stable and that were recorded in at least three trials per experimental condition were included in the analyses (a minimum of ~100 trials, typically ~350 trials).

#### 5. Data analysis

##### 5.1. Mean firing rate of neurons

Mean firing rates were calculated separately for each frequency and for *left-* and *right-starting* metronome trials using a 50 ms window displaced at 10 ms steps (Fig. 2). To establish each neuron’s preferred location (Fig. 3a), we computed the cross-correlogram between the firing rate and stimulus location. A positive correlation index at lag 0 indicated a *left-starting* preference. We used this index to group trials into preferred and non-preferred starting location. Firing rates of each neuron were normalized (*z*-values) by subtracting the mean and dividing by the standard deviation across trials.

##### 5.2. Time-frequency analyses

Time-frequency power plots (spectrograms) were calculated with Matlab R2019a (The Mathworks, Natick, MA, USA) with the function *spectrogram*, using a 500 ms Hamming window displaced at 50 ms steps. Essentially equal results were obtained using multi-taper spectral analysis with the Chronux Toolbox^3^. Spectrograms were calculated separately for *left-* and *right-starting* metronomes and then subtracted from each other to illustrate the dynamic changes in power associated with the metronome’s tempo (Fig. 3b). Mean power (white lines in Fig. 3b) was calculated by averaging across frequencies. The power for each recording session was normalized (*z*-values) before averaging.

##### 5.3. Prediction of behavioral choices using support-vector machines (SVM)

To test whether the firing rates of single neurons on single trials contained information about the monkeys’ decision, we trained a linear SVM to classify trials into *leftwards* or *rightwards* choices. We included correct and incorrect trials (Extended Data Fig. 2b). To have sufficiently long trials, we selected trials with go-cues between 3-6 intervals (3-4 for the SMA; we did no use trials with only 1 or 2 elapsed intervals). The three metronome tempos (500, 750, 1000 ms) were pooled by using a common time axis (in interval units), and a different classifier was used for each go-cue. We performed a 10-fold validation procedure, and a single sided *t-test* was used to determine whether the activity of each neuron significantly predicted the behavioral choices (Fig. 3c).

##### 5.4. Principal Component Analysis (PCA) and demixed-PCA (dPCA)

To visualize how population activity evolves as a function of time, we used Principal Component Analysis to obtain the first 3 PCs from the mean firing rate of all neurons over each task condition. A single PC transformation was applied to the complete data matrix of each recorded area. Each row of the data matrix contained the mean activity of a single neuron, and each column contained the firing rate of the neurons as a function of time, with task conditions concatenated in the time dimension (columns). In Extended Data Fig. 3a-b we examined the first 2 PCs as a function of time, conditioned by spatial location and tempo. In main Fig. 4 we examined the evolution of population activity (dynamics of the state space) by plotting combinations of PC1, PC2, and PC3. The trajectories across these state spaces were used to illustrate examples of brain areas with selectivity for the metronome’s location and tempo. To estimate the number of PCs needed to account for 80% of activity variance (mean ± s.d.; Fig. 5d) we used a bootstrapping procedure in which we took 500 samples of 130 randomly selected neurons, with replacement, from each area. Sampling the same number of neurons allowed to control for possible confounds introduced by the different number of recorded neurons in each area.

To precisely measure the extent to which each cortical area represents (1) the spatial location of the metronome (left or right), (2) the speed of the metronome (interval duration: 500, 750, 1000 ms), and (3) total elapsed time since the beginning of the trial, we made use of a dimensionality reduction technique that, in addition to capturing activity variance (like PCA), it quantifies how much this activity represents tasks parameters (Figure 5a-c; Extended Data Figs. 4–5). The capacity of each neuron to encode the metronome’s characteristics (quantified by dPCS weights) allowed us to establish the contribution of each cortical area in solving the metronome task. As in standard PCA, the dPCA weights are calculated from the concatenated activity of all the recorded neurons across areas and across conditions ^4^.

